# Modulating Innate Immune Responses to Curli Fibers Through Protein Engineering

**DOI:** 10.64898/2026.03.23.713613

**Authors:** Shanna Bonanno, Rutvi Sheta, Teena Ramu, Shriya Verenkar, Darren Kim, Eva Bessette, Petit Pierre, Neel S. Joshi

**Affiliations:** Department of Bioengineering, Northeastern University, Boston, MA 02115; Department of Chemistry and Chemical Biology, Northeastern University, Boston, MA 02115; Department of Biology, Northeastern University, Boston, MA 02115; Department of Chemistry, Tufts University, Medford, MA 02155

**Keywords:** curli fibers, *Escherichia coli* Nissle 1917, Toll-like receptors, innate immunity, microbial amyloids, immune modulation, pattern recognition receptors, engineered living materials, mucosal immunity

## Abstract

Curli fibers produced by *Escherichia coli* are functional amyloids that activate Toll-like receptor 2 (TLR2), initiating innate immune responses at mucosal surfaces. While microbiome-derived curli contribute to host-microbe interactions, their intrinsic immunostimulatory activity limits their utility as programmable scaffolds for engineered probiotic systems, and dysregulated TLR2 activation has been linked to inflammatory bowel disease, systemic lupus erythematosus, neurodegeneration, and sepsis. Here, we engineered *E. coli* Nissle 1917 to produce modified curli fibers designed to inhibit TLR2 through two mechanistically distinct strategies: steric shielding via silk-elastin-like protein sequences, and direct receptor antagonism via a known TLR2 antagonist, staphylococcal superantigen-like protein 3 (SSL3). Both engineered variants assembled into structurally intact amyloid fibers and exhibited significantly reduced intrinsic TLR2-dependent NF-κB activation in reporter cells. In competitive inhibition assays against structurally diverse TLR2 agonists, the SSL3 fusion achieved near-complete inhibition maintained under rising agonist load, while steric shielding provided moderate, agonist-class-dependent inhibition. In primary human monocyte-derived dendritic cells, the SSL3 fusion robustly attenuated IL-8 secretion and transcriptional induction of IL-8, IL-6, and IL-1β, whereas steric shielding produced only partial attenuation that did not translate to broad inflammatory suppression. These results establish engineered curli as a tunable platform for receptor-specific modulation of innate immune signaling and highlight the broader potential of modular microbial amyloids as programmable interfaces for engineering host-microbe interactions at mucosal surfaces.

**IMPORTANCE:** Bacteria residing in the gut produce protein fibers called curli that potently activate the immune system through a receptor called Toll-like receptor 2 (TLR2). While TLR2 plays a beneficial role in maintaining gut health, its overactivation drives chronic inflammation in conditions including inflammatory bowel disease, autoimmune diseases, neurodegenerative diseases, and sepsis, and curli fibers have been directly implicated in several of these conditions. Here, we engineered curli fibers produced by the probiotic *E. coli* Nissle 1917 to inhibit TLR2 activation, transforming a naturally inflammatory bacterial fiber into a programmable immune modulator. We demonstrated that direct receptor antagonism, rather than steric shielding, is required for effective immune modulation in primary human immune cells, establishing a design principle for engineering bacteria-derived fibers as programmable interfaces with host immunity. The modularity of the curli scaffold positions this platform as a broader tool for programming interactions between probiotic bacteria and the mucosal immune system.

## INTRODUCTION

The gastrointestinal tract harbors a dense microbial community that plays a central role in shaping mucosal immunity (1–3). Intestinal epithelial and immune cells continuously sample microbial components, maintaining a dynamic balance between tolerance to commensal organisms and protective responses against pathogens (4, 5), a balance that pattern recognition receptors (PRRs) mediate through detection of conserved microbial structures and initiate innate immune signaling (6–8).

Toll-like receptor 2 (TLR2) is a key sensor of bacterial products at mucosal surfaces, including lipoproteins, peptidoglycan, lipoteichoic acid, and zymosan (9, 10). TLR2 forms heterodimers with either TLR1 or TLR6 to recognize structurally diverse ligands (11) and signals through the MyD88-dependent pathway to activate NF-κB and downstream pro-inflammatory cytokine transcription (12–14). While TLR2 contributes to mucosal homeostasis, including reinforcement of epithelial barrier integrity (9, 15), its dysregulation is implicated in inflammatory bowel disease (IBD), systemic lupus erythematosus (SLE), neurodegenerative diseases, and sepsis (13, 16–23). In active IBD, TLR2 activation contributes to a self-reinforcing inflammatory cycle in which tissue damage releases endogenous damage-associated molecular patterns (DAMPs) that further activate TLR2 and NF-κB signaling, perpetuating chronic mucosal inflammation (13, 16, 17). Current therapeutic strategies rely largely on systemic immunosuppression or broadly acting biologics, which carry risks of off-target immunosuppression and fail to address the localized, receptor-level dysregulation. New approaches are needed that can selectively and locally tune TLR2-mediated responses at mucosal surfaces.

Among the microbial structures sensed by TLR2 are curli fibers, functional amyloids produced by *Escherichia coli* and other Enterobacteriaceae (24). Composed primarily of CsgA, which polymerizes into β-sheet-rich fibrils, curli contribute to biofilm formation, host colonization, and cell-cell signaling (25), and engage the TLR2/TLR1 heterodimer to promote pro-inflammatory cytokine production (26, 27). Microbiome-derived curli are directly implicated in broader inflammatory pathologies beyond the gut: curli-DNA composites formed during enteric biofilm formation drive autoantibody production in lupus-prone mice, linking gut biofilms to SLE pathogenesis (19). Studies have implicated curli cross-seeding with α-synuclein in the enteric nervous in Parkinson’s disease progression (28, 29). These findings suggest that locally modulating curli-TLR2 interactions could have therapeutic relevance across a range of pathologies involving TLR2.

Curli fibers are also uniquely attractive as programmable scaffolds for engineered probiotic systems. Unlike soluble protein therapeutics, curli self-assemble extracellularly at the mucosal interface, enabling localized, matrix-associated presentation of functional domains. This scaffold-immobilized format may prolong receptor contact time and sustain local ligand presentation under the continuous flow and clearance conditions of the intestinal lumen, in ways that freely diffusing molecules cannot replicate (30). CsgA is highly amenable to genetic modification, allowing incorporation of diverse peptide or protein domains without disrupting fiber assembly (31). Prior work has demonstrated the utility of both native and engineered curli in a range of mucosal disease-relevant contexts (32–36), and *E. coli* Nissle 1917 (EcN) has been used as a delivery vehicle for curli-based living therapeutics given its genetic tractability and history of use in the gut (37–41). However, the intrinsic ability of native curli to activate TLR2 presents a fundamental challenge for their use as immunomodulatory scaffolds, requiring strategies that retain curli’s structural advantages while redirecting their immunological properties.

Here, we hypothesized that engineering EcN to produce curli fibers that inhibit TLR2 activation would enable selective modulation of innate immune responses at mucosal surfaces. We pursued two mechanistically distinct strategies: steric shielding of TLR2-interacting surfaces using silk-elastin-like protein (SELP) sequences, and direct receptor antagonism through staphylococcal superantigen-like protein 3 (SSL3), a characterized TLR2-blocking protein. We validated production, assembly, and structural integrity of modified curli fibers, evaluated TLR2-dependent signaling in reporter cells, and assessed competitive inhibition against diverse TLR2 agonists. Furthermore, we assessed inflammatory responses in primary human monocyte-derived dendritic cells (MoDCs).

Because curli-producing commensals contribute to TLR2-driven pathology through direct receptor activation, an engineered EcN producing TLR2-antagonistic curli could compete with endogenous curli in the gut niche, reducing the immunostimulatory burden at its source. More broadly, the modularity of the CsgA scaffold suggests this approach could serve as a foundation for programming interactions between probiotic bacteria and host PRRs in a receptor-specific manner. Together, this work establishes a genetically encoded strategy for modulating innate immune signaling through engineered microbial amyloids and provides a mechanistic framework for programming host-microbe receptor interactions at mucosal surfaces.

## RESULTS

### Design and Engineering of Curli Variants to Modulate TLR2 Recognition

Native curli fibers are potent activators of the TLR2/TLR1 heterodimer (Fig. 1A) (24, 26). To generate curli fibers capable of inhibiting TLR2 activation (Fig. 1B), we exploited the modularity of the CsgA scaffold to introduce C-terminal fusion domains representing two complementary strategies: steric shielding via fusion of flexible, biocompatible SELP sequences to physically occlude TLR2-interacting surfaces on the fiber, and direct receptor antagonism via fusion of the SSL3 protein, which occupies the TLR2 binding site and blocks downstream activation (42, 43). We screened a library of SELP variants for TLR2 signaling activity, and the S6E11 sequence was selected as the lead steric shielding candidate based on its ability to reduce NF-κB reporter activation. This design produced a panel of three engineered EcN strains expressing CsgA wild-type (WT), CsgA-S6E11, and CsgA-SSL3 for evaluating how distinct modes of CsgA modification influence TLR2-dependent innate immune signaling. Relevant plasmid maps and corresponding DNA and protein sequences are provided in the supplementary materials (Fig. S1, Table S1-2).

**Figure 1.**
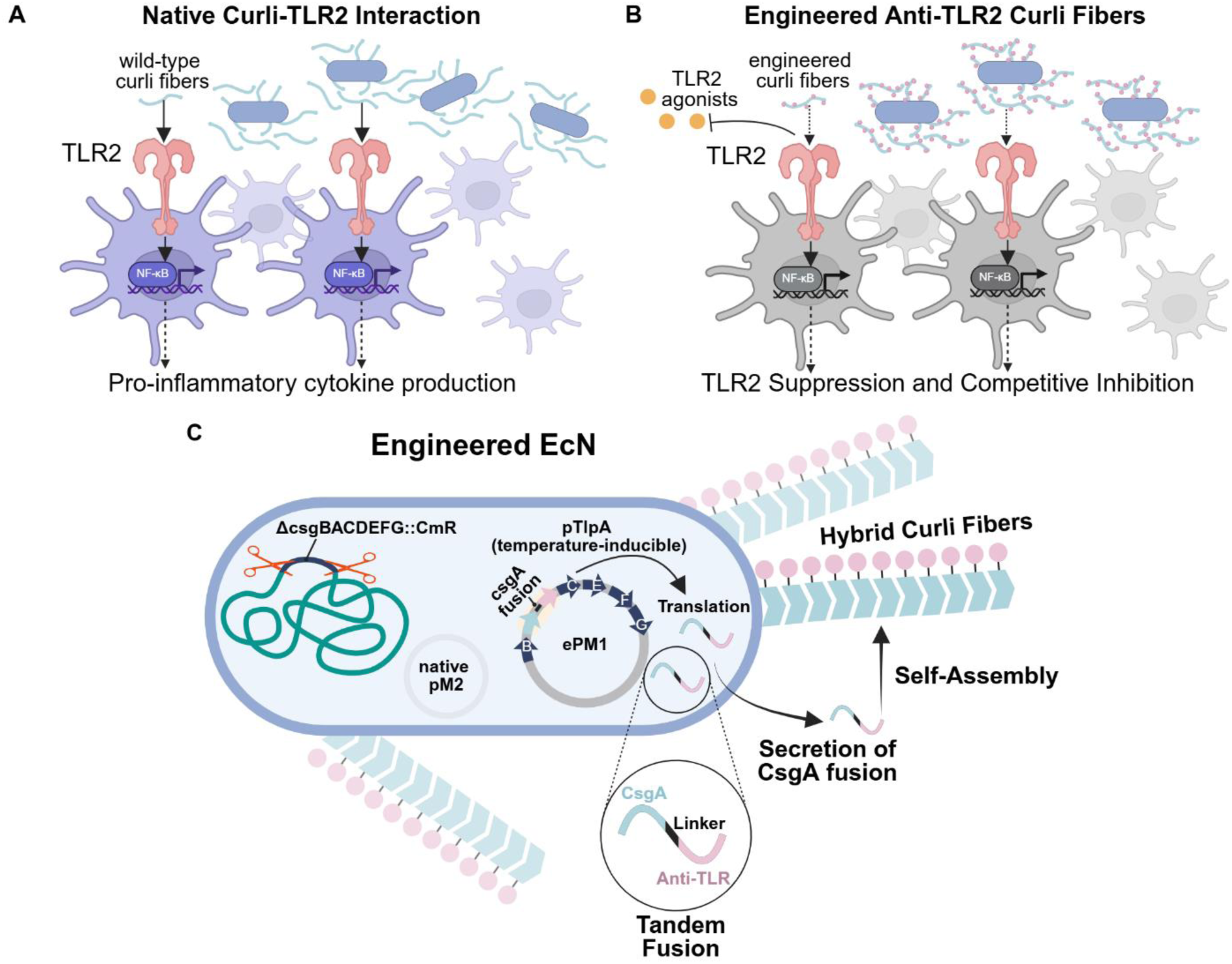
Conceptual Overview of Curli Fiber Engineering to Modulate TLR2 Signaling. **(A)** Wild-type curli fibers produced by *E. coli* activate host TLR2/TLR1, triggering NF-κB signaling and pro-inflammatory responses (24, 26, 44). **(B)** Engineered curli fibers exhibit reduced intrinsic TLR2 activation and inhibit TLR2 signaling induced by diverse agonists, providing a tunable platform for selective modulation of innate immune responses. **(C)** Engineering strategy. The endogenous curli operon was deleted from EcN (Δ*csgBACEFG::cmR*) to eliminate native fiber production (32). A temperature-inducible synthetic curli operon encoding secretion machinery (*csgBCEFG*), along with engineered *csgA* variants, was introduced on an engineered pMUT1-derived plasmid (ePM1) (45, 46). Engineered variants consisted of C-terminal fusions of functional domains to CsgA via flexible glycine-serine linker containing internal 6×His and E-tag affinity tags. Upon thermal induction, EcN secretes modified CsgA monomers that self-assemble extracellularly into hybrid curli fibers.

All studies were performed in EcN, a clinically validated probiotic chassis capable of colonizing the gut mucosa and producing genetically encoded biomaterials *in situ* (47). To ensure that all immune phenotypes could be attributed exclusively to engineered constructs, endogenous curli operon components were deleted from the chromosome (EcN Δ*csgBACDEFG::cmR*), eliminating background fiber production (Fig. 1C) (32). Engineered CsgA variants were expressed from a synthetic curli operon, co-transcribed with the secretion and assembly genes *csgB, csgC, csgE, csgF,* and *csgG* (25), and cloned into an engineered pMUT1-derived plasmid (ePM1) (46). The ePM1 backbone was selected because EcN naturally maintains the pMUT1 cryptic plasmid, enabling development of a compatible vector with the potential for antibiotic-free maintenance relevant to future therapeutic deployment. Expression was placed under control of the temperature-sensitive *pTlpA* promoter, which enables inducible curli production in response to physiological temperature cues and limits constitutive expression that could impose metabolic burden on the host bacterium (45). Experiments were performed in EcN strains lacking endogenous pMUT1 to ensure controlled, exclusive expression from the ePM1 plasmid.

### Engineered Curli Variants Retain Amyloid Formation and Fiber Assembly

Because reduced immune activation could arise from impaired fiber production rather than altered receptor recognition, we sought to confirm that all constructs produced extracellular amyloid fibers comparable to wild-type curli before proceeding to immune characterization.

Whole-cell Congo red binding assays demonstrated robust amyloid production across engineered strains, with signal normalized to cell density (OD_600_) indicating efficient fiber formation comparable to wild-type constructs (Fig. 2A). Consistent with extracellular display of engineered monomers, whole-cell ELISA using anti-E-tag detection confirmed surface-accessible fusion domains across all variants (Fig. 2B). Immunoblot analysis of cell lysates further verified expression of full-length CsgA fusion proteins at the expected molecular weights, confirming intact fusion protein production for each construct (Fig. 2C).

**Figure 2.**
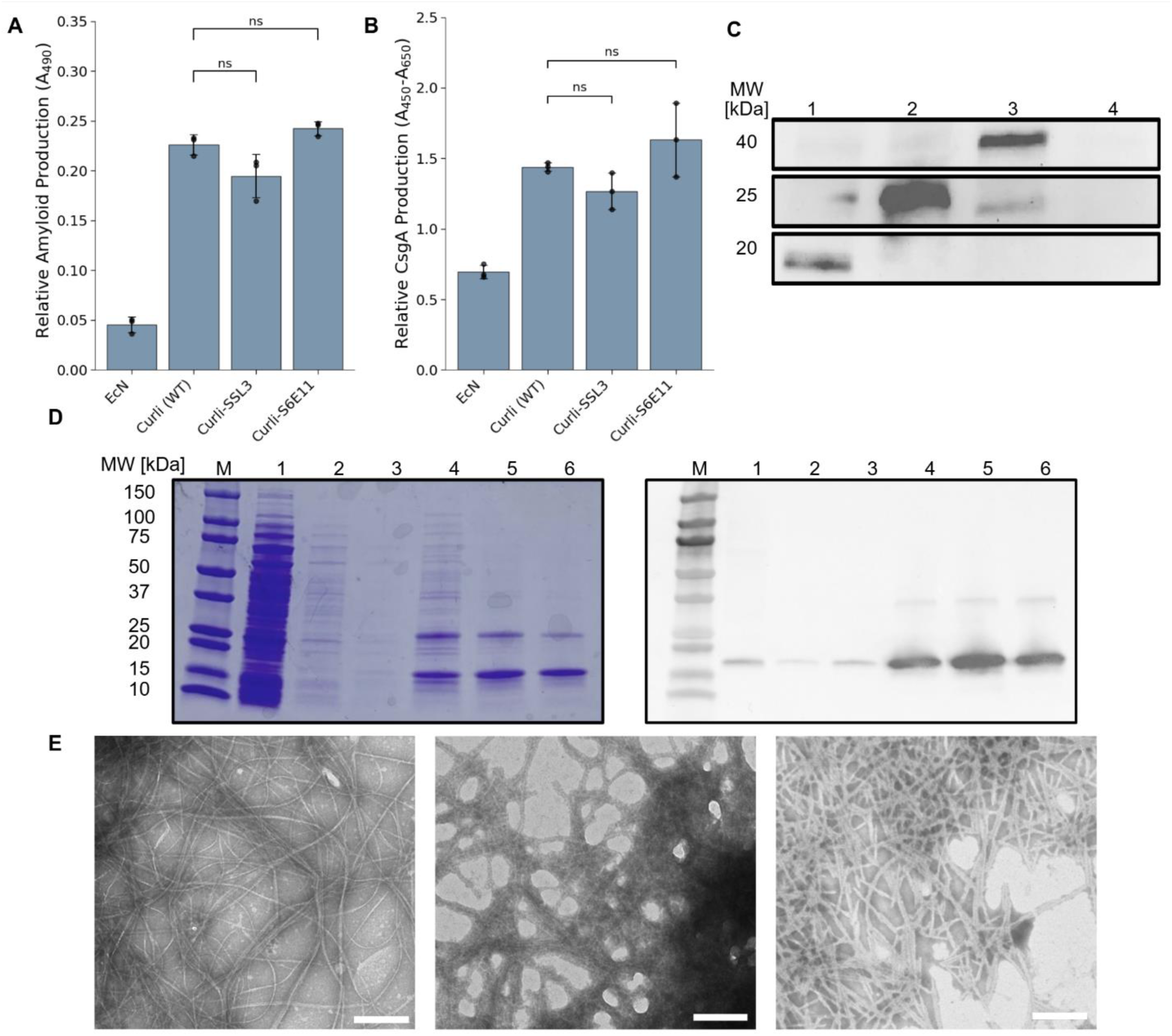
Production and Characterization of Curli Fiber Variants from Engineered EcN. **(A)** Congo red binding assay of induced cell cultures, normalized to OD_600_, demonstrating comparable amyloid production across engineered strains. **(B)** Whole-cell ELISA measuring surface-accessible CsgA fusion domains using anti-E-tag antibody, normalized to OD_600_, confirming extracellular display of fusion domains. **(A-B)** The EcN condition represents the EcN Δ*csgBACEFG::cmR* untransformed control. Statistical comparisons between wild-type curli and each fusion were performed using Welch’s t-tests (ns, not significant). **(C)** Western blot analysis of whole-cell lysates confirming expression of full-length CsgA fusion proteins at expected molecular weights. Lanes: (1) Curli (WT), (2) Curli-S6E11, (3) Curli-SSL3, (4) EcN Δ*csgBACEFG::cmR* untransformed control. **(D)** Representative SDS-PAGE and anti-6×His Western blot of purified wild-type curli fibers confirming recovery of intact His-tagged CsgA monomers. Left: Coomassie-stained SDS-PAGE; Right: Anti-6×His Western blot. Lanes: (M) protein marker, (1) lysate flow-through, (2) 50 mM phosphate buffer wash (KPi buffer), (3) 12.5 mM imidazole wash, (4–6) 125 mM imidazole elution fractions. Purification data for engineered fusion variants are provided in Fig. S3. **(E)** Transmission electron microscopy of purified curli fibers showing dense fibrillar assemblies characteristic of amyloid morphology across all variants. Left: Curli (WT); Middle: Curli-SSL3; Right: Curli-S6E11. Scale bars, 200 nm.

To enable dose-controlled biochemical and immunological characterization independent of bacterial background effects, curli variants were also expressed and purified as isolated fibers. Initial purification from standard BL21(DE3) yielded protein samples with residual lipopolysaccharide (LPS) contamination that drove elevated inflammatory responses in primary human MoDCs, confounding curli-specific immune readouts (Fig. S2). We therefore expressed constructs in the endotoxin-free ClearColi™ BL21(DE3) strain using isopropyl β-d-1-thiogalactopyranoside (IPTG)-inducible vectors, minimizing LPS contamination and ensuring that downstream immune phenotypes could be attributed specifically to curli fibers rather than residual endotoxin. We purified curli assemblies under denaturing conditions using 8 M guanidine hydrochloride. SDS-PAGE and anti-His immunoblotting confirmed successful recovery of intact His-tagged CsgA monomers (Fig. 2D), with consistent recovery across all engineered variants (Fig. S3). After purification, TLR4 activation in HEK-Blue™ hTLR4 reporter cells was low and comparable across all ClearColi™-purified fiber preparations, confirming a marked reduction in LPS contamination relative to BL21(DE3)-purified samples (Fig. S4).

To confirm that denaturing purification conditions did not compromise fusion domain integrity, we assessed secondary structure retention of refolded SSL3 by circular dichroism spectroscopy. SSL3 was selected as the representative structured domain for this analysis, as SELP sequences are intrinsically disordered and insoluble curli fibers are not amenable to CD measurement. Refolded SSL3 exhibited spectra consistent with its native beta-sheet rich secondary structure, with dominant peaks at 210-220 nanometers (nm) (negative) and 190 nm (positive) (Fig. S5), suggesting that guanidinium treatment does not irreversibly disrupt domain folding and supporting the functional integrity of fusion domains following purification (48).

Transmission electron microscopy of purified fibers confirmed extracellular fibrillar assemblies with dense, β-sheet-rich morphology characteristic of canonical amyloid architecture across all three variants (Fig. 2E). Fiber morphology, density, and overall structure appeared comparable between wild-type and engineered constructs, indicating that C-terminal domain fusions do not disrupt the amyloid assembly process.

Together, these results demonstrate that engineered curli variants are efficiently produced, assemble into structurally intact amyloid fibers and retain domain stability following purification. The comparable fiber morphology and production levels across variants establish that downstream differences in immune activation arise from engineered alterations to receptor interactions rather than defects in fiber formation or surface display.

### Engineered Curli Variants Exhibit Reduced TLR2-Dependent Activation in HEK Reporter Cells

Having confirmed that engineered curli fibers assemble into structurally intact amyloid fibers, we next evaluated how these modifications influence TLR2-dependent signaling using HEK-Blue™ hTLR2 reporter cells. These cells express the human TLR2/TLR1 heterodimer and a secreted alkaline phosphatase (SEAP) reporter under NF-κB control, enabling quantitative measurement of receptor activation (Fig. 3A). Prior studies indicate that curli fibers specifically engage this TLR2/TLR1 heterodimer, establishing it as the primary mediator of NF-κB activation in this system (26).

**Figure 3.**
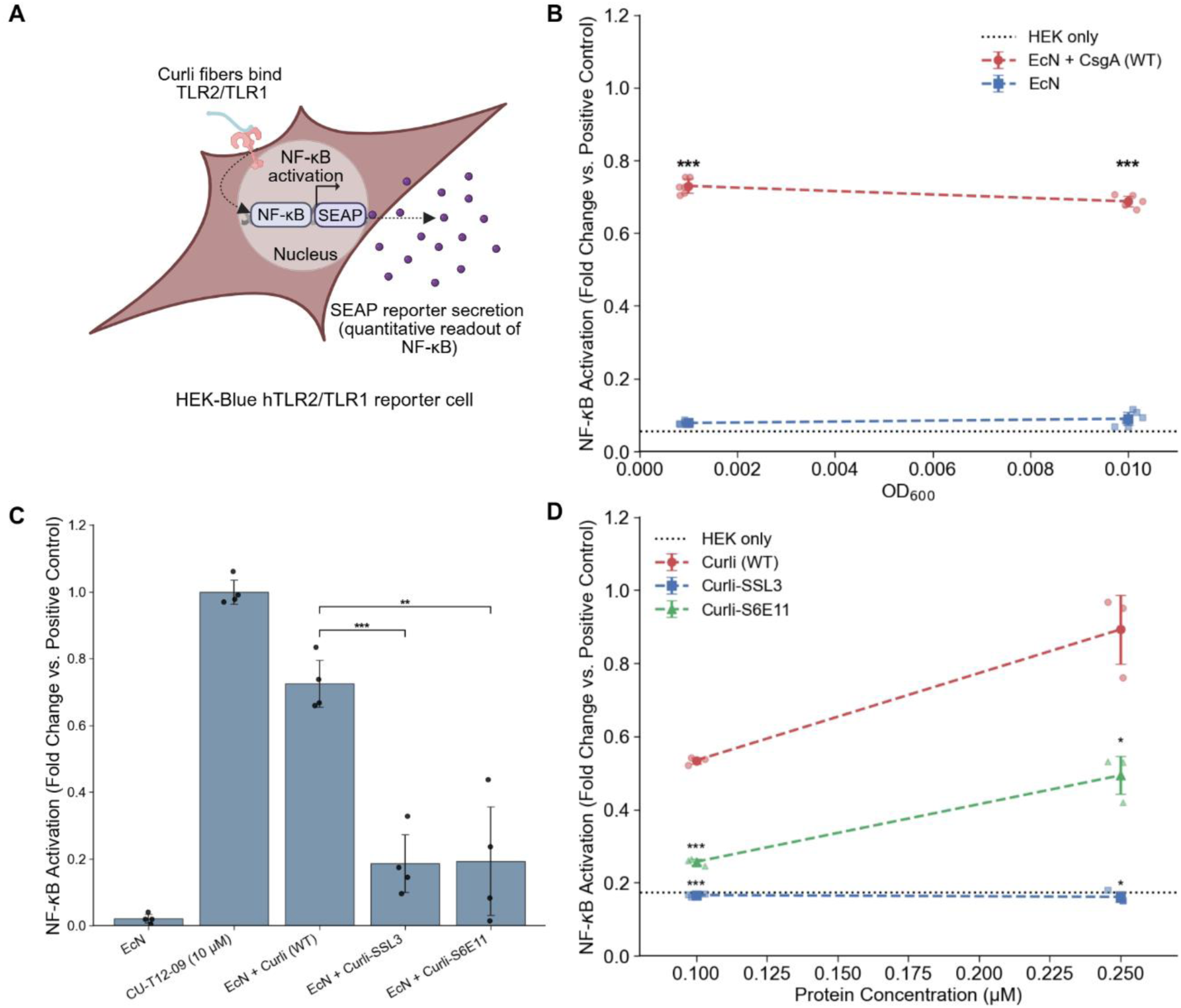
TLR2-dependent responses to engineered curli variants in HEK-Blue™ reporter cells. **(A)** Schematic of the HEK-Blue™ hTLR2/TLR1 SEAP reporter system. HEK cells expressing the TLR2/TLR1 heterodimer produce SEAP under NF-κB control upon receptor activation, enabling quantitative measurement of TLR2-dependent signaling. **(B)** NF-κB activation in HEK-TLR2 cells treated with curli-producing EcN or curli-deficient EcN (Δ*csgBACEFG::cmR*) across two bacterial densities (OD600 = 0.001 and 0.01). The dotted black horizontal line indicates the no-treatment negative control baseline. Statistical analyses were performed using two-way ANOVA (protein × concentration interaction), followed by planned two-sided Welch’s t-tests with Bonferroni correction for multiple comparisons between EcN + curli (WT) and EcN (no curli control) at each concentration. **(C)** NF-κB activation in HEK-TLR2 cells treated with curli-deficient EcN, CU-T12-9, or EcN expressing wild-type or engineered curli variants (Curli-SSL3, Curli-S6E11) at OD600 = 0.01 (multiplicity of infection, MOI ≈ 160). Pairwise comparisons between wild-type curli and each engineered variant were performed using Welch’s t-test with Bonferroni correction for multiple comparisons. **(D)** Dose-dependent NF-κB activation in HEK-TLR2 cells treated with purified Curli (WT), Curli-SSL3, or Curli-S6E11 fibers at 0.1 and 0.25 µM. The dotted black horizontal line indicates the no-treatment negative control baseline. Statistical analyses were performed using two-way ANOVA (protein × concentration interaction), followed by planned two-sided Welch’s t-tests with Bonferroni correction for multiple comparisons between engineered curli variants and Curli (WT) at each concentration. **(B-D)** Reporter activity is presented as fold change relative to CU-T12-9. Data represent mean ± standard deviation (SD) with individual replicates shown. Statistical significance is indicated as *, p < 0.05; **, p < 0.01; ***, p < 0.001; ns, not significant.

We first established that TLR2 activation in this system is curli-dependent. HEK-TLR2 cells were treated with either curli-producing EcN or curli-deficient EcN (Δ*csgBACEFG::cmR*) across two bacterial densities, with the synthetic TLR2/TLR1 agonist CU-T12-9 serving as a positive control. Curli-deficient EcN produced minimal reporter activation at both densities, indistinguishable from untreated cells, whereas wild-type curli-producing EcN induced robust NF-κB signaling, reaching 12.9-fold and 12.1-fold activation at OD600 = 0.001 and 0.01, respectively, relative to the untreated control (Fig. 3B). This confirms that curli fiber production, rather than other bacterial components, is necessary and sufficient for TLR2 pathway activation in this system.

We next compared the effect of induced EcN cells producing either wild-type curli or engineered variants on HEK-TLR2 response after normalizing bacterial density (OD600 = 0.01; MOI ≈ 160). Wild-type curli induced NF-κB activation at approximately 0.73-fold relative to CU-T12-9, while both engineered variants produced significantly lower responses, with Curli-SSL3 and Curli-S6E11 at approximately 0.19-fold relative to CU-T12-9 (Fig. 3C).

To confirm receptor specificity and rule out non-immune confounding factors, we performed two additional controls. First, identical treatments in parental HEK-Blue™ Null1 cells, which share the same genetic background as HEK-TLR2 cells but lack TLR2 expression, produced no detectable NF-κB activation across any curli-producing strain (Fig. S6), confirming that signaling is TLR2-dependent and not due to nonspecific cellular responses. Second, cell viability assessed across all treatment conditions was comparable to untreated controls (Fig. S7), confirming that differences in reporter activation reflect modulation of receptor signaling rather than cytotoxic effects.

To confirm these findings under conditions that isolate fiber-intrinsic receptor engagement from bacterial background, we treated HEK-TLR2 cells with purified curli fibers normalized by protein mass, providing the most direct measure of how engineered modifications alter TLR2 activation per unit of fiber. Wild-type curli induced dose-dependent NF-κB activation, reaching approximately 0.53-fold and 0.89-fold of the CU-T12-9 positive control at 0.1 µM and 0.25 µM, respectively (Fig. 3D). Curli-SSL3 maintained consistently low activation across both doses (approximately 0.16-fold of at both concentrations), and these reductions were statistically significant relative to wild-type at both concentrations. Similarly, Curli-S6E11 produced significantly lower activation than Curli (WT) at 0.1 µM (approximately 0.26-fold) and at 0.25 µM (approximately 0.49-fold).

Finally, to confirm that observed phenotypes were not caused by proteolytic cleavage of fusion domains during cell exposure, curli fibers were recovered after overnight incubation with HEK cells, depolymerized, and analyzed by Western blot. All variants retained full-length CsgA fusions at expected molecular weights (Fig. S8), confirming that fusion domains remain intact and fiber-associated throughout the assay and that reduced signaling reflects engineered modulation of TLR2 engagement rather than domain loss.

Collectively, these results establish that both engineered curli variants exhibit significantly reduced intrinsic TLR2/TLR1 activation in a curli-dependent, receptor-specific, and concentration-resolved manner.

### Engineered Curli Variants Inhibit TLR2 Activation by Diverse Agonists

Having established that engineered curli variants exhibit reduced intrinsic TLR2 activation, we next asked whether these fibers could actively modulate receptor signaling by inhibiting activation induced by external TLR2 agonists. To test this, HEK-Blue™ hTLR2 reporter cells were pre-incubated with wild-type or engineered curli fibers prior to stimulation with defined TLR2 ligands representing structurally distinct classes: CU-T12-9, a synthetic small-molecule TLR2/TLR1 agonist; heat-killed *S. typhimurium* (HKST), a complex pathogen-derived stimulus; and lipoteichoic acid (LTA), a gram-positive bacterial cell wall component (Fig. 4A).

**Figure 4.**
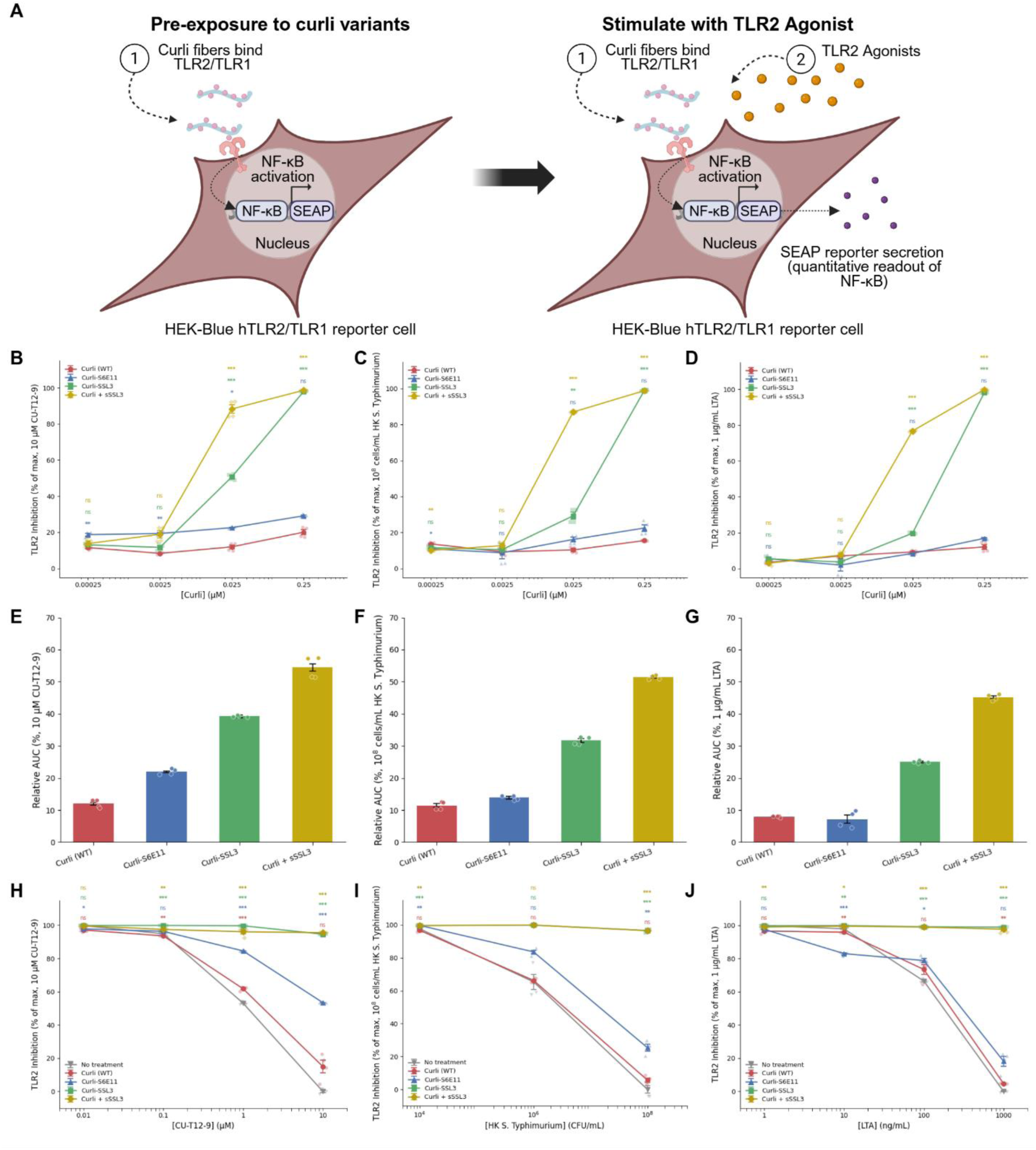
Engineered curli variants inhibit TLR2 activation across diverse agonists. **(A)** Schematic of competition assays. HEK-Blue™ hTLR2 reporter cells were pre-incubated with wild-type or engineered curli fibers before stimulation with defined TLR2 agonists. **(B-D)** Percent inhibition of NF-κB activation as a function of curli antagonist concentration (0.00025-0.25 µM, log scale), with agonist concentration held constant: **(B)** 10 µM CU-T12-9, **(C)** 10⁸ cells/mL HKST, **(D)** 1 µg/mL LTA. Inhibition is normalized to agonist-alone stimulation (0% inhibition) and untreated HEK cells (100% inhibition). **(E-G)** Relative area under the curve (AUC) derived from individual replicate antagonist titration curves (panels B-D; n = 4 per construct) for each construct across all three agonists: **(E)** 10 µM CU-T12-9, **(F)** 10⁸ cells/mL HKST, **(G)** 1 µg/mL LTA. **(H-J)** Percent inhibition of NF-κB activation as a function of agonist concentration (log scale), with curli concentration held constant at 0.25 µM: **(H)** CU-T12-9 (0 µM-10 µM), **(I)** HKST (0-10⁸ cells/mL), **(J)** LTA (0-1 µg/mL). Inhibition is normalized to the highest dose of agonist-alone stimulation (0% inhibition) and untreated HEK cells (100% inhibition). Statistical comparisons in panels B-D were performed using Welch’s t-tests vs. Curli (WT) with Bonferroni correction for multiple pairwise comparisons. Statistical comparisons in panels H-J were performed using Welch’s t-test vs. No treatment with Bonferroni correction for multiple pairwise comparisons. Data in panels B-J represent mean ± standard error of the mean (SEM) of four independent replicates. Statistical significance is indicated as *, p < 0.05; **, p < 0.01; ***, p < 0.001; ns, not significant.

In antagonist titration experiments, cells were stimulated with a fixed agonist concentration: 10 µM CU-T12-9, 10⁸ cells/mL HKST, or 1 µg/mL LTA, while curli concentrations were varied across four doses (0.00025-0.25 µM) (Fig. 4B-D). At the highest curli concentration tested (0.25 µM), Curli-SSL3 and Curli (WT) supplemented with equimolar soluble SSL3 (Curli (WT) + sSSL3) both achieved near-complete inhibition, reaching >98% inhibition across all three agonists. In contrast, Curli-S6E11 achieved only moderate inhibition (17-29%) and Curli (WT) remained largely ineffective (12-20%) across all agonists at 0.25 µM.

Because the antagonist titration experiments sampled a limited number of curli concentrations, sigmoidal curve fitting was not reliably applicable to all conditions, and IC_50_ values could not be robustly estimated. We therefore used the relative area under the antagonist titration curve (Relative AUC) as a proxy for overall inhibitory activity across the tested concentration range. Relative AUC was calculated for each individual replicate curve and expressed as percentage of theoretical maximum (100% inhibition across all concentrations tested), allowing direct comparison across agonists and constructs (Fig. 4E-G).

Across all three agonists, Curli + sSSL3 consistently achieved the highest relative AUC, reflecting broad and potent inhibition across the full concentration range tested: 54.45 ± 1.08% for CU-T12-9, 51.40 ± 0.32% for HKST, and 45.18 ± 0.44% for LTA. Curli-SSL3 showed consistently strong inhibitory activity across all three agonists (CU-T12-9: 39.24 ± 0.41%; HKST: 31.67 ± 0.63%; LTA: 25.01 ± 0.22%), substantially exceeding both Curli-S6E11 and Curli (WT). Curli-S6E11 showed modest inhibitory activity for CU-T12-9 (21.94 ± 0.27%) but minimal activity against HKST (13.86 ± 0.39%) and LTA (7.15 ± 1.30%), suggesting agonist-dependent differences in its inhibitory mechanism. Curli (WT) showed the lowest relative AUC across all conditions (CU-T12-9: 12.02 ± 0.45%; HKST: 11.39 ± 0.61%; LTA: 7.98 ± 0.15%), consistent with its minimal antagonist activity observed in the titration curves.

To complement the antagonist titration experiments, we next examined inhibitory activity under conditions of fixed antagonist and variable agonist concentration, holding curli at 0.25 μM while titrating agonist across a log-scale range. Curli-SSL3 and Curli (WT) + sSSL3 maintained >95% inhibition across the full agonist dose range for all three agonists tested (Fig. 4H-J), demonstrating that SSL3-mediated TLR2 blockade is robust to increasing agonist load. Curli (WT) showed minimal inhibitory activity (<15%) across all agonist concentrations, consistent with its inability to compete effectively with agonist binding at the TLR2 receptor. Curli-S6E11 showed variable and agonist-dependent performance: approximately 53% inhibition at the highest CU-T12-9 concentration (Fig. 4H), but only approximately 25% and 16% at the highest concentrations of HKST (Fig. 4I) and LTA (Fig. 4J).

Together, these results establish that engineered curli fibers incorporating the SSL3 antagonist domain function as potent modulators of TLR2 signaling across structurally diverse agonists, maintaining near-complete inhibition even under rising agonist load. The consistency of this effect across a synthetic agonist, a complex pathogen-derived stimulus, and a bacterial cell wall component demonstrates broad-spectrum activity at the receptor level.

### Engineered Curli Variants Attenuate TLR2-Dependent Inflammatory Responses in Primary Human MoDCs

To determine whether the immunomodulatory effects observed in reporter assays translated to primary immune cells, we evaluated inflammatory responses in human MoDCs. Dendritic cells were selected because they serve as key antigen-presenting cells at mucosal interfaces and central regulators of immune activation and tolerance (49, 50). We restricted experiments to purified curli fiber stimulation to isolate TLR2-dependent responses from confounding bacterial background signals.

Primary human CD14⁺/CD16⁻ classical monocytes were isolated from a leukopak by immunomagnetic enrichment and differentiated into MoDCs using established cytokine-driven protocols (Fig. 5A) (51).

**Figure 5.**
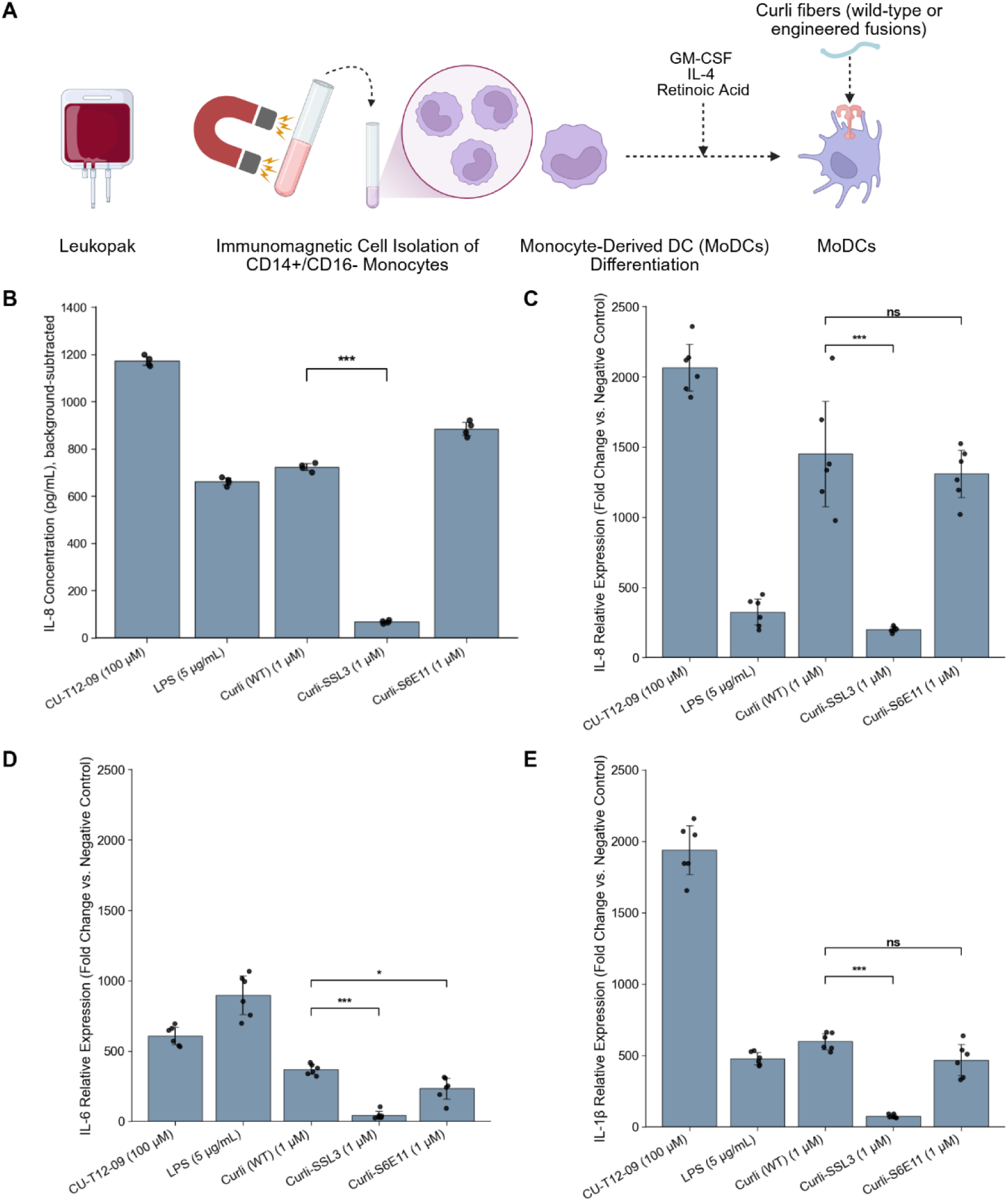
Engineered curli variants attenuate TLR2-dependent inflammatory responses in primary human MoDCs. **(A)** Schematic of MoDC generation. CD14⁺/CD16⁻ classical monocytes were isolated from a leukopak by immunomagnetic enrichment and differentiated into MoDCs using established cytokine-driven protocols prior to stimulation with purified curli fibers (51). **(B)** IL-8 secretion measured by ELISA following 24 h exposure to LPS (the canonical MoDC maturation stimulus), CU-T12-9 (TLR2/TLR1 positive control), Curli (WT), Curli-SSL3, or Curli-S6E11. Values are reported in pg/mL with background subtracted (untreated MoDC baseline set to 0). Statistical comparison between Curli (WT) and Curli-SSL3 was performed using Welch’s t-test. **(C-E)** RT-qPCR analysis of inflammatory gene expression following 24 h curli exposure: **(C)** IL-8, **(D)** IL-6, **(E)** IL-1β. Data are expressed as fold change relative to untreated MoDCs (set to 1). Statistical comparisons between Curli (WT) and each engineered variant were performed using Welch’s t-tests with Bonferroni correction for multiple pairwise comparisons. Data in panels B-E represent mean ± SD. Statistical significance is indicated as *, p < 0.05; **, p < 0.01; ***, p < 0.001; ns, not significant.

Exposure to purified Curli (WT) induced robust IL-8 secretion (724 pg/mL: above background), reaching 62% of the response induced by CU-T12-9 (Fig. 5B), consistent with prior reporter assay results. Curli-SSL3 produced significantly lower IL-8 levels (71 pg/mL), demonstrating effective attenuation of inflammatory activation. Curli-S6E11 did not reduce IL-8 secretion relative to Curli (WT) and trended toward slightly higher cytokine production (898 pg/mL), indicating an inability to antagonize TLR2 in the receptor environment of primary dendritic cells.

Transcriptional responses were further characterized by RT-qPCR quantification of canonical inflammatory genes IL-8, IL-6, and IL-1β, expressed as fold change relative to untreated MoDCs (Fig. 5C-E). Consistent with ELISA measurements, Curli (WT) strongly upregulated all three inflammatory transcripts, while Curli-SSL3 significantly reduced gene induction across all markers tested (7-fold reduction for IL-8, 8-fold reduction for IL-6, 8-fold reduction for IL-1β vs. Curli (WT)). IL-8 and IL-1β levels in Curli-S6E11-treated cells were not significantly different from Curli (WT); however, IL-6 transcripts were modestly but significantly reduced (1.6-fold), suggesting partial, cytokine-specific transcriptional attenuation that does not extend to broad suppression of inflammatory response.

To confirm that differences in inflammatory signaling were not attributable to cytotoxic effects, MoDC viability was assessed by flow cytometry using live/dead staining following curli exposure. All treatment conditions maintained high cell viability relative to untreated controls (Fig. S9), confirming that reduced cytokine production by Curli-SSL3 reflects modulation of immune signaling rather than loss of viable cells.

To confirm that the observed inflammatory responses were TLR2-dependent, MoDCs were pretreated with a small-molecule inhibitor of TLR2 (TL2-C29) prior to curli stimulation. Blockade by TL2-C29 reduced Curli (WT)-induced IL-8, IL-6, and IL-1β production by approximately 85%, 80%, and 84%, respectively (Fig. S10), confirming that the inflammatory response to curli in MoDCs is TLR2-mediated.

Together, these results demonstrate that incorporation of the SSL3 antagonist domain into curli fibers is sufficient to attenuate TLR2-dependent inflammatory activation in primary human immune cells, an effect confirmed at both the protein and transcriptional level.

## DISCUSSION

This study demonstrated that genetically engineered curli fibers can be repurposed from inherently immunostimulatory microbial amyloids into programmable modulators of innate immune signaling. By modifying the CsgA scaffold to alter TLR2 receptor engagement, we established a genetically encoded strategy for tuning TLR2-dependent responses while preserving amyloid assembly and extracellular fiber formation.

A central finding is that while both the S6E11 SELP fusion and the SSL3 fusion reduced intrinsic TLR2-dependent activation by curli, only the SSL3 fusion attenuated inflammatory responses robustly across experimental systems. Curli-S6E11 reduced TLR2 activation in simplified reporter systems, but this attenuation did not translate to effective blockade of exogenous TLR2 agonists or to broad suppression of inflammatory responses in primary MoDCs. While the S6E11 sequence was selected as the lead SELP candidate from an initial library screen, SELP sequences represent a broad and tunable class of intrinsically disordered polymers whose steric and physiochemical properties can be systematically varied through changes in sequence length, composition, and repeat architecture, which may yield variants with enhanced shielding efficacy (52, 53). The partial attenuation observed with S6E11 in certain contexts suggests that steric shielding retains promise as a design strategy and warrants further exploration, even if specific receptor antagonism proved more effective in the present study.

Direct receptor antagonism by SSL3 provides a more absolute inhibitory mechanism that remained effective regardless of cellular context, agonist class, or agonist concentration. Competition experiments established that Curli-SSL3 functions as a competitive modulator, and the robustness of SSL3-mediated inhibition across three structurally distinct agonist classes and under rising agonist load supports its suitability for deployment in complex, ligand-rich inflammatory environments.

Notably, Curli (WT) + sSSL3 achieved modestly higher AUC than Curli-SSL3 at submaximal curli concentrations, most evident at 0.025 µM, while both converged to comparable inhibition at 0.25 µM. This indicated that fiber immobilization does not enhance intrinsic antagonist potency under static *in vitro* conditions. However, these experiments were performed under static cell culture conditions that do not recapitulate the dynamic flow environment of the gut mucosa. *In vivo*, tethering SSL3 to a self-assembling scaffold produced directly at the mucosal surface could sustain local antagonist presentation and prolong receptor contact time under the continuous fluid flow and clearance conditions of the intestinal lumen in ways that freely diffusing soluble proteins cannot (54, 55). The advantage of the fiber-tethered format therefore likely lies in its biological delivery context rather than intrinsic binding potency, and future *in vivo* studies will be needed to evaluate whether scaffold immobilization confers a functional advantage under physiologically relevant conditions.

The success of the SSL3 fusion also points toward a broader design opportunity. Having established that direct antagonism of TLR2 via a fiber-displayed protein domain is both feasible and effective, we propose that the modularity of the CsgA scaffold could support analogous fusions targeting other innate immune receptors. Targeting innate immune receptors is an emerging therapeutic strategy with broad relevance across inflammatory, autoimmune, and infectious disease contexts (38, 56, 57), and curli fibers, which are amenable to diverse C-terminal fusions without loss of assembly, are well-positioned as a scaffold for this class of intervention. Looking further ahead, combinations of steric and antagonistic domains on the same fiber, or libraries of CsgA fusions targeting distinct receptor-ligand interfaces, could enable programmable, receptor-specific tuning of mucosal immune responses in a genetically encoded format.

These findings are directly relevant to the dysregulation of TLR2 signaling in chronic inflammatory disease. In active IBD, TLR2 is upregulated on lamina propria immune cells and sustained activation by DAMPs and commensal-derived ligands perpetuates a self-reinforcing inflammatory cycle that current treatments act downstream of rather than at the receptor level (13, 16, 17). A probiotic engineered to produce TLR2-antagonistic curli directly at the mucosal surface represents a mechanistically distinct approach targeting receptor-level dysregulation at its site of origin rather than suppressing downstream effectors systemically. A particular advantage of the fiber-tethered format is that EcN colonizes the same mucosal niche as endogenous curli-producing commensals and other TLR2 agonists, positioning an engineered strain to continuously present TLR2 antagonist at the site of agonist production (32, 58). Furthermore, curli have been implicated in broader TLR2-driven pathologies including autoantibody production in lupus-prone mice (19), and α-synuclein cross-seeding in the enteric nervous system linked to Parkinson’s disease progression (28, 29), suggesting that locally attenuating curli-TLR2 interactions could have relevance across multiple disease contexts. Whether Curli-SSL3 can reduce TLR2-driven inflammation *in vivo* remains to be established, and future studies in relevant disease models will be necessary to evaluate therapeutic feasibility.

In summary, this work establishes that the immunological properties of a bacterial extracellular matrix component can be rationally redesigned without compromising structural function. By transforming curli into a competitive TLR2 modulator, we demonstrated that synthetic biology can reprogram the innate immune signaling identity of a microbial amyloid, establishing curli as a programmable interface between engineered probiotic bacteria and host mucosal immunity.

## MATERIALS AND METHODS

### Bacterial Strains, Plasmids, and Genetic Engineering

All plasmids used in this study were constructed using standard molecular biology techniques. DNA fragments were amplified using Q5 High-Fidelity DNA Polymerase Master Mix (New England Biolabs). Custom oligonucleotides were ordered from Genewiz (Azenta Life Sciences) and custom gene blocks were ordered from Twist Bioscience. Gibson Assembly (New England Biolabs) was used to assemble DNA fragments for plasmid construction and site-directed mutagenesis (New England Biolabs) was used to introduce sequence modifications where required.

All plasmids were transformed into One Shot Mach1 Phage-Resistant Chemically Competent *E. coli* (Thermo Fisher Scientific) and plated onto LB agar with appropriate antibiotics. Isolated colonies were inoculated into LB broth and cultured overnight at 30°C for temperature-sensitive *pTlpA* constructs or 37°C otherwise, with shaking at 225 RPM. Plasmids were purified using the Qiagen MiniPrep kit and verified by whole-plasmid sequencing (Plasmidsaurus).

The following vectors were used in this study: pMUT1 and pET21d(+). The pMUT1 vector was isolated from EcN and genes were inserted as previously described (45). The SELP S6E11 sequence was obtained from a plasmid deposited on Addgene: pMAL-c5X-SELP0K was a gift from Ashutosh Chilkoti (Addgene plasmid # 67007; http://n2t.net/addgene:67007; RRID: Addgene_67007). Constructs used in this study include: pMUT1-pTlpA-csgAwt, pMUT1-pTlpA-csgA-S6E11, pMUT1-pTlpA-csgA-SSL3, pET21d-csgAwt, pET21d-csgA-S6E11, pET21d-csgA-SSL3, and pET21d-SSL3. Plasmid maps and corresponding DNA and protein sequences for all constructs used in this study are provided in supplementary materials (Fig. S1, Table S1-2). Plasmids generated in this study have been deposited at Addgene and will be publicly available upon publication.

### Bacterial Culture and Curli Expression

EcN Δ*csgBACEFG* ΔpMUT1, a derivative of EcN described previously (46), was used for all whole-cell and bacterial co-culture experiments. ClearColi™ BL21(DE3) (Lucigen) was used for protein purification. Both strains were transformed with the corresponding plasmids by electroporation. Transformants were plated onto LB agar containing 100 µg/mL carbenicillin (Teknova) and incubated overnight at 30°C for temperature-sensitive *pTlpA* constructs or 37°C otherwise.

Single colonies were inoculated into 5 mL LB broth containing 100 µg/mL carbenicillin and grown overnight at 30°C or 37°C with shaking at 225 RPM. For *pTlpA*-driven expression in EcN, overnight cultures were diluted 1:1000 into fresh LB containing 100 µg/mL carbenicillin and incubated overnight at 37°C with shaking to induce curli expression via temperature upshift. For pET21d-based expression in ClearColi™, overnight cultures were similarly diluted 1:1000 into fresh LB containing 100 µg/mL carbenicillin, induced with 100 µM IPTG, and incubated overnight at 37°C with shaking.

### Congo Red binding assay

One milliliter of bacterial culture was pelleted at 5,500 × g for 10 minutes and the supernatant was discarded. The cell pellet was resuspended in 1 mL of 0.025 mM Congo red and incubated on a rotator for 10 minutes at room temperature. Cells were pelleted at 16,500 × g for 10 minutes, and the supernatant was aliquoted in triplicate into a 96-well plate. Absorbance was measured at 490 nm using a microplate reader (Molecular Devices, SpectraMax M5 Multi-Mode Microplate Reader). Normalized curli production was calculated by subtracting the absorbance of a blank (0.025 mM Congo red without cells) from the sample absorbance and normalizing to the OD600 of the original culture.

### Whole cell filtration ELISA

Bacterial cultures were normalized to an OD600 of 0.5 in TBS, and 200 µL of each normalized culture was pipetted in triplicate into a 96-well filter plate (MilliporeSigma Multiscreen®, 0.22 µm). Cultures were filtered through the membrane under vacuum, and wells were washed three times with 300 µL TBST with vacuum applied between each wash.

Wells were blocked with 200 µL of blocking solution (1% BSA, 0.01% hydrogen peroxide in TBST) for 1.5 hours at 37°C, filtered through, and washed three times with 300 µL TBST. Fifty microliters of HRP-conjugated anti-E-tag antibody (NB600-526, 1:5,000 in 1% BSA/TBST) was added per well and incubated for 1.5 hours at room temperature, followed by four washes with 200 µL TBST.

Signal was developed by adding 100 µL 3,3′,5,5′-tetramethylbenzidine (TMB) ELISA reagent per well and incubating at room temperature in the dark until the desired signal was reached. The reaction was stopped with 50 µL of 2 M sulfuric acid, and 100 µL from each well was transferred to a clear 96-well plate. Absorbance was measured at 450 nm with background subtraction at 650 nm.

### Curli purification

Following overnight induction of cytosolic curli fibers, cultures were pelleted at 10,000 × g for 20 minutes and stored at –80°C. Pellets were resuspended in 8 M guanidine hydrochloride in 50 mM potassium phosphate (KPi) buffer (pH 7.2) at 0.06× the original culture volume. Samples were incubated at room temperature for 1 hour with gentle rotation, followed by sonication (60% amplitude, 20 sec on/off, three cycles) at room temperature. Samples were then centrifuged at 10,000 × g for 20 minutes at 4°C, and the supernatant was transferred to a fresh tube. A second round of sonication (20 sec on/off, three cycles) was performed.

HIS-Select® HF Nickel Affinity Gel (MilliporeSigma, cat. #H0537) was added at 0.03× the supernatant volume, and the mixture was incubated for 1 hour at room temperature with gentle rotation. The resin was loaded onto a column and washed with four column volumes of sterile water, followed by four column volumes of 8 M guanidine hydrochloride in 50 mM KPi buffer (pH 7.2). The column was drained under compressed air and washed with two column volumes of 50 mM KPi buffer (pH 7.2). Three milliliters of 12.5 mM imidazole in KPi buffer (pH 7.2) was added and drained under compressed air. The column was then capped at the bottom, and 3 mL of 125 mM imidazole in 50 mM KPi buffer (pH 7.2) was added and incubated for 3-5 minutes before elution into three fractions.

To prevent polymerization, protein analysis was performed immediately following purification. Samples were desalted and buffer-exchanged into 50 mM KPi buffer (pH 7.2) using Zeba Spin Desalting Columns (7K MWCO, cat. #89892, Thermo Scientific). Purity and identity were assessed by SDS-PAGE with Coomassie staining and Western blot, and protein concentration was determined by BCA assay, as described below.

#### SDS-PAGE

For whole-cell experiments, curli fibers were secreted and self-assembled during induction. To depolymerize curli fibers, the desired volume of cell culture (normalized to OD_600_) was pelleted by centrifugation at 16,000 × g for 3 minutes. Pellets were resuspended in 100% hexafluoroisopropanol (HFIP) at half the original culture volume. HFIP was evaporated under vacuum using a Savant SpeedVac (Thermo Fisher Scientific) at 65°C for 30-60 minutes. Samples were then resuspended in SDS loading buffer (Laemmli buffer containing 0.1 M dithiothreitol (DTT)).

For purified curli samples, SDS loading buffer (Laemmli buffer containing 0.1 M DTT) was added directly following purification while fibers remained in a denatured state. All samples were heated at 95°C for 10 minutes, then loaded onto a 4-10% Mini-PROTEAN TGX gradient gel (Bio-Rad) alongside Precision Plus Protein Dual Color Standards (Bio-Rad). Electrophoresis was performed in Tris-glycine-SDS running buffer at 90 V for 15 minutes to stack proteins, followed by 130 V until the dye front reached the bottom of the gel.

#### Coomassie staining

Following SDS-PAGE, gels were stained with 0.1% (w/v) Coomassie Brillian Blue R-250 in 50% methanol and 10% glacial acetic acid for 1 hour at room temperature with gentle shaking. Gels were then detained overnight in 40% methanol and 10% glacial acetic acid at room temperature with shaking until background was sufficiently cleared, before imaging using a Bio-Rad ChemiDoc MP imaging system.

#### Western blot

Following SDS-PAGE, proteins were transferred to a polyvinylidene fluoride (PVDF) membrane using the Invitrogen™ iBlot™ 2 Dry Blotting System with the P0 transfer program. The membrane was blocked for 1 hour at room temperature in TBST containing 5% (w/v) bovine serum albumin (BSA) with gentle shaking, followed by overnight incubation at 4°C with gently shaking in horseradish peroxidase (HRP)-conjugated 6×His Tag monoclonal antibody (MA1-80218, Invitrogen) diluted 1:2,500 in 1% (w/v) BSA in TBST. The membrane was then washed three times for 5 minutes each in TBST with gentle shaking. Signal was developed using Pierce™ ECL Western Blotting Substrate (cat. #32209, Thermo Fisher Scientific) and images using a Bio-Rad ChemiDoc MP imaging system.

#### BCA assay

Protein concentration was determined using the Pierce™ Dilution-Free™ Rapid Gold BCA Protein Assay Kit (Thermo Fisher Scientific, cat. #A55861) according to the manufacturer’s instructions. Samples were combined with working reagent and incubated at room temperature for 5 minutes. Absorbance was measured at 480 nm using a microplate reader (Molecular Devices, Spectramax M5 Multi-Mode Microplate Reader). Protein concentrations were calculated by interpolation from a pre-diluted BSA standard curve (50-1000 μg/mL).

#### Transmission Electron Microscopy

Purified curli fiber samples were prepared for transmission electron microscopy by negative staining. 200-mesh formvar-carbon grids (Electron Microscopy Sciences) were held with tweezers, and a small droplet of vortexed protein sample was applied to the grid surface and incubated for 1 minute to allow adsorption. Excess liquid was gently wicked away using filter paper. Grids were then stained with 2% phosphotungstic acid (pH 7.0) for 1 minute, and excess stain was removed by wicking. Grids were allowed to air-dry completely at room temperature prior to imaging. Samples were imaged using a JEM-101 transmission electron microscope (JEOL ltd.) operated at 60 kV and equipped with an AMT XR-41B CCD camera (2048 × 2048 pixels) for digital image acquisition.

#### Circular Dichroism

Purified protein samples were buffer-exchanged into 50 mM KPi buffer (pH 7.2) and normalized to a final concentration of 0.2 mg/mL. Samples were loaded into a 1 mm pathlength quartz cuvette (Starna Cells, Inc. Atascadero, CA USA) that had been pre-rinsed with KPi buffer (pH 7.2). Circular dichroism spectra were collected on a J-1500 Circular Dichroism Spectrophotometer (JASCO Inc., Easton, MD, USA) at 25°C. A blank with KPi buffer solution was measured prior to samples acquisition and automatically subtracted from all spectra before data export.

Spectra were recorded from 260-190 nm using a bandwidth of 1.0 nm, data intervale of 0.1 nm, scanning speed of 50 nm/min, detector integration time of 2 s, and nitrogen purge flow rate at 8 L/min. Five accumulations were averaged for each spectrum. Raw CD signal in millidegrees (mdeg) was converted using Jasco Spectra Manager software and reported as molar ellipticity ([θ]) using Equation 1 below where θ is the ellipticity (mdeg), c is the protein concentration in mol/L, l is the cell pathlength (cm), and R is the number of amino acid residues in the protein (59).

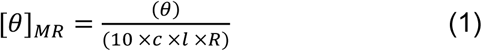

#### Endotoxin Assay

Endotoxin levels in purified fiber preparations were quantified using the Pierce™ Chromogenic Endotoxin Quant Kit (Thermo Fisher Scientific, cat. #A39552S) according to the manufacturer’s instructions. Protein samples and endotoxin standards were prepared in endotoxin-free water and added to endotoxin-free microplate wells. Following addition of LAL reagent and incubation at 37°C, the reaction was stopped with the supplied stop solution and absorbance was measured at 405 nm. Endotoxin concentrations were calculated from a standard curve and reported as endotoxin units per milliliter (EU/mL). All samples were measured in technical replicates.

#### HEK-Blue™ Cell Lines

HEK-Blue™ hTLR2 reporter cells (hkb-htlr2, InvivoGen) are derivatives of HEK293 cells that stably express human TLR2 and an NF-κB-inducible secreted alkaline phosphatase (SEAP) reporter. Cells were maintained in Dulbecco’s Modified Eagle Medium (DMEM; 4.5 g/L glucose, 2 mM L-glutamine) supplemented with 10% (v/v) fetal bovine serum (FBS), penicillin-streptomycin (100 U/mL–100 µg/mL), 100 µg/mL Normocin™, and 1× HEK-Blue™ Selection according to the manufacturer’s instructions. HEK-Blue™ Null1 cells (hkb-null1, InvivoGen) served as the parental TLR2-negative control and were maintained in the same medium supplemented with 1× Zeocin in place of HEK-Blue™ Selection.

#### HEK-Blue™ Reporter Assays

HEK-Blue™ reporter cells were seeded at 5.6 × 10⁴ cells per well in 96-well plates and allowed to adhere overnight in growth medium lacking selection antibiotics. Prior to co-culture, EcN strains were washed three times by centrifugation and resuspension in sterile PBS to remove residual media components. Cells were then stimulated with washed EcN strains at OD600 = 0.01 (MOI ≈ 160) or purified curli fibers at 0.25 µM unless otherwise specified, for 18 hours at 37°C in a humidified incubator with 5% CO₂.

#### SEAP Reporter Assay for NF-κB Activation

NF-κB activation was quantified using QUANTI-Blue™ colorimetric detection reagent (InvivoGen) to measure SEAP activity in cell culture supernatants. Following stimulation, supernatants were transferred to a fresh 96-well plate containing QUANTI-Blue™ reagent and incubated at 37°C until color development. Absorbance was measured at 655 nm using a microplate reader (Molecular Devices, SpectraMax M5 Multi-Mode Microplate Reader). Background signal from unstimulated controls was subtracted, and SEAP activity was normalized to the CU-T12-9 positive control, assigned a value of 1.

#### LDH Cell Viability

Cell viability following stimulation was assessed using the LDH-Blue™ Cytotoxicity Assay Kit (InvivoGen, cat. #rep-ldh-1) according to the manufacturer’s instructions. Following the stimulation period, culture supernatants were collected and transferred to a 96-well plate. Supernatants were mixed with LDH-Blue™ reagent solution at a 1:1 ratio and incubated at room temperature for 30 minutes in the dark. Absorbance was measured at 650 nm using a microplate reader (Molecular Devices, SpectraMax M5 Multi-Mode Microplate Reader).

Percent cytotoxicity was calculated relative to untreated cells (spontaneous LDH release) and cells treated with lysis buffer (maximum LDH release) using Equation 2 below.

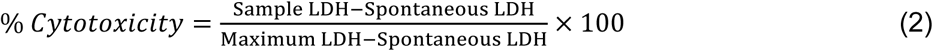

#### SSL3 Purification

Soluble His-tagged SSL3 protein was expressed in ClearColi™ BL21(DE3) (Lucigen) from a pET-based expression vector. Overnight starter cultures were grown in LB broth with 100 µg/mL carbenicillin at 37°C with shaking at 250 rpm. Expression cultures were inoculated at a 1:100 dilution and grown at 37°C with shaking overnight, induced by addition of IPTG to a final concentration of 100 µM. Cells were harvested by centrifugation at 4,000 × g for 20 min at 4°C and pellets were stored at −80°C until lysis.

Cell pellets were resuspended in BugBuster^®^ Protein Extraction Reagent (MilliporeSigma, cat. #70584-M) at 5 mL per gram of wet cell mass, supplemented with Benzonase nuclease (25 U/mL) and protease inhibitor cocktail. Resuspensions were incubated at room temperature for 20 min with rocking, followed by clarification at 16,000 × g for 20 min at 4°C. The clarified lysate was applied to a Ni-NTA resin column (HIS-Select® HF Nickel Affinity Gel, MilliporeSigma, cat. #H0537) pre-equilibrated with binding buffer (50 mM sodium phosphate, 300 mM NaCl, 20 mM imidazole, pH 8.0). The column was washed twice with 5-10 mL of wash buffer (50 mM sodium phosphate, 300 mM NaCl, 20 mM imidazole, pH 8.0). Protein was eluted in six 2 mL fractions at stepwise increasing imidazole concentrations: two fractions at 20 mM, two at 200 mM, and two at 500 mM imidazole in 50 mM sodium phosphate, 300 mM NaCl, pH 8.0. Peak fractions were identified by SDS-PAGE and pooled. Imidazole was removed by buffer exchange into 50 mM potassium phosphate, pH 7.2, using Zeba Spin Desalting columns (7K MWCO, Thermo Scientific). Protein concentration was determined by BCA assay as described above, and purity was confirmed by SDS-PAGE with Coomassie staining and anti-6×His Western blot. Purified protein was aliquoted and stored at −80°C.

#### Monocyte Isolation from Human Leukopak

Human monocytes were isolated from a leukopak (STEMCELL Technologies) using the EasySep™ Human Monocyte Isolation Kit (STEMCELL Technologies, cat. #19359) according to the manufacturer’s instructions. Briefly, leukopak samples were mixed thoroughly, and an aliquot was diluted 1:10 in a 1:100 solution of acetic acid and methylene blue for cell counting using a hemocytometer. The remaining leukopak was transferred to a sterile 50 mL conical tube, washed with EasySep™ Buffer (PBS, 2% FBS, 1 mM EDTA), and pelleted at 300 × g for 10 min at room temperature.

Red blood cells were lysed by resuspending the pellet in ammonium chloride solution and incubating on ice for 5-7 min, followed by pelleting at 300 × g. Cells were washed twice with EasySep™ Buffer to remove platelets and counted. Cell concentration was adjusted to 5 × 10⁷ cells/mL for magnetic isolation. The monocyte isolation cocktail (50 µL/mL) and magnetic particles (50 µL/mL) were added sequentially with incubation on ice for 10 min each, followed by magnetic separation using the EasySep™ magnet. The enriched monocyte fraction was collected, and monocyte purity and yield were assessed by trypan blue exclusion.

Isolated monocytes were pelleted and resuspended in FBS containing 10% dimethyl sulfoxide (DMSO) at approximately 4 × 10⁶ cells/mL, frozen overnight at −80°C in a controlled-rate freezing container (Mr. Frosty, Thermo Fisher Scientific), and subsequently stored in liquid nitrogen until use.

#### Primary Human MoDC Differentiation

Frozen human monocytes were thawed rapidly in a 37°C water bath, washed twice with PBS containing 2% FBS and 0.5 mM EDTA, and counted using a hemocytometer with trypan blue exclusion to assess viability. Cells were resuspended in DC differentiation medium consisting of RPMI 1640 supplemented with 1× GlutaMAX, 1% penicillin–streptomycin, 50 ng/mL granulocyte-macrophage colony-stimulating factor (GM-CSF), 35 ng/mL interleukin-4 (IL-4), and 10 nM retinoic acid, as previously described (51). Cells were seeded at 5 × 10⁵ cells per well in 24-well tissue culture-treated plates. Cultures were maintained at 37°C with 5% CO₂. Fresh differentiation medium (0.2 mL per well) was added every other day without removing existing medium. On day 6, fully differentiated immature MoDCs were harvested for downstream stimulation assays.

#### MoDC Stimulation Assays

Immature MoDCs were generated as described above and stimulated on day 6 of differentiation. Purified curli fiber variants were diluted to 1 µM in fresh DC differentiation medium and added to cultures for 48 h at 37°C with 5% CO₂. Positive controls included LPS (5 µg/mL) the canonical MoDC maturation stimulus and CU-T12-9 (100 µM) for TLR2/TLR1-dependent activation. Following the 48 h incubation, supernatants were collected for cytokine quantification by ELISA, and cells were harvested for RNA extraction and flow cytometric viability assessment as described in the respective sections below.

#### TLR2 Receptor Blocking Assay

To confirm that curli fiber-induced inflammatory responses were mediated primarily through TLR2 signaling, MoDCs were pretreated on day 6 of differentiation with the TLR2 signaling inhibitor TL2-C29 (100 μM; Invivogen, cat. #inh-c29) for 3 hours at 37°C. Without removing the inhibitor, cells were then stimulated with either purified curli fibers (1 µM) or CU-T12-9 (100 µM) and incubated for 48 hours at 37°C with 5% CO₂. Parallel control conditions included purified curli fibers and CU-T12-9 without TL2-C29 pretreatment. Following incubation, cells were harvested for RNA extraction and proinflammatory cytokine expression (IL-8, IL-6, IL-1β) was assessed as described below.

#### IL-8

IL-8 secretion from MoDCs was measured using the LEGEND MAX™ Human IL-8 ELISA Kit (BioLegend, cat. 431504) following the manufacturer’s instructions. Briefly, 96-well high-binding ELISA plates were coated with capture antibody and incubated overnight at 4°C. Plates were washed three times with wash buffer and blocked with assay buffer for 1 hour at room temperature. Cell culture supernatants or IL-8 standards were added at 100 µL per well and incubated at room temperature for 2 hours. Wells were washed three times before addition of detection antibody and incubated for 1 hour at room temperature. After three additional washes, avidin-HRP reagent was added and incubated for 30 minutes. Following a final wash, substrate solution was added and allowed to develop for 15–20 minutes at room temperature in the dark. The reaction was stopped with stop solution, and absorbance was measured at 450 nm using a microplate reader (Molecular Devices, SpectraMax M5 Multi-Mode Microplate Reader), with 570 nm used for background subtraction. IL-8 concentrations were calculated from a standard curve generated using the supplied IL-8 standards. All samples were measured in technical triplicates, and mean values were reported. Background IL-8 from untreated MoDCs was subtracted from all conditions prior to reporting.

#### RNA Isolation

Total RNA was extracted from MoDCs using the Quick-RNA Microprep Kit (Zymo Research, cat. # R1050) according to the manufacturer’s instructions. Cells were lysed directly in RNA Lysis Buffer, and an equal volume of 95–100% ethanol was added to the lysate and mixed thoroughly. The lysate was transferred to a Zymo-Spin IC column and centrifuged at 12,000 × g for 30 seconds, and the flow-through was discarded. The column was washed with 400 µL RNA Wash Buffer. Genomic DNA was removed on-column by adding a mixture of 5 µL DNase I (1 U/µL) and 35 µL DNA Digestion Buffer directly to the column matrix and incubating for 15 minutes at room temperature. The column was then washed sequentially with 400 µL RNA Prep Buffer, 700 µL RNA Wash Buffer, and a final wash with 400 µL RNA Wash Buffer. RNA was eluted in 15 µL DNase/RNase-free water. RNA concentration and purity were assessed using a NanoDrop spectrophotometer (Thermo Fisher Scientific), with acceptable samples defined by a 260/280 ratio of ∼2.0 and a 260/230 ratio >1.8.

#### RT-qPCR

Relative gene expression was assessed by one-step reverse transcription quantitative PCR (RT-qPCR) using the Luna® Universal One-Step RT-qPCR Kit with SYBR Green (New England Biolabs) according to the manufacturer’s instructions. Gene-specific primers for IL-8, IL-6, IL-1β, and GAPDH were designed using NCBI Primer-BLAST with consistent parameters to enable simultaneous analysis (Supplementary Table 3). RNA samples were thawed on ice and diluted to a working concentration of 10 ng/µL. Master mixes were prepared containing SYBR Green RT-qPCR reagents and forward and reverse primers at a final concentration of 0.2 µM each. One microliter of RNA template was added per 20 µL reaction, and all reactions were performed in 96-well PCR plates sealed with adhesive plate seals. Plates were briefly centrifuged before cycling. RT-qPCR was performed on a Bio-Rad CFX96 Real-Time PCR System using the following thermal cycling conditions: reverse transcription at 55°C for 10 min, initial denaturation at 95°C for 1 min, followed by 40 cycles of 95°C for 10 s and 60°C for 30 s. Melt curve analysis was performed after amplification to confirm primer specificity. Relative gene expression was calculated using the ΔΔCt method with GAPDH as the housekeeping reference gene.

#### Flow Cytometry

Following stimulation, MoDCs were harvested by collecting culture supernatants and detaching adherent cells with Accutase (0.5 mL per well, 1-10 min at 37°C). Non-adherent and adherent cell fractions were combined, pelleted at 500 × g for 5 min at 4°C, and washed twice with cold fluorescence-activated cell sorting (FACS) buffer (PBS, 2% FBS, 1 mM EDTA). Cell counts were determined, and 5 × 10⁵ cells were aliquoted per sample. Fc receptors were blocked using Human TruStain FcX (BioLegend, 5 µL per sample, 10 min at room temperature). Cell viability was assessed using LIVE/DEAD™ Fixable Aqua Dead Cell Stain (Invitrogen, Cat. No. L34957), incubated in 100 µL PBS with 1 µL dye for 30 min at room temperature in the dark. After staining, cells were washed twice with FACS buffer, resuspended in 200 µL FACS buffer, and analyzed immediately on a Beckman Coulter CytoFLEX S Cell Analyzer (Institute for Chemical Imaging of Living Systems, Northeastern University). Unstained and single-stain compensation controls were prepared using AbC™ and ArC™ bead kits (Invitrogen) according to the manufacturer’s instructions. Viable cells were identified as negative for the amine-reactive Live/Dead dye, and data were analyzed using FlowJo software.

#### Statistics and reproducibility

Data are presented as mean ± standard deviation (SD) with individual replicates shown. A significance threshold of p < 0.05 was applied for all tests, indicated as *, p < 0.05; **, p < 0.01; ***, p < 0.001; ns, not significant.

Welch’s t-test was used for pairwise comparisons between two groups with unequal variances. Where multiple pairwise comparisons were made from the same dataset, Bonferroni correction was applied, including comparisons between wild-type and engineered curli variants in whole-cell reporter assays (Fig. 3C), competitive inhibition assays (Fig. 4B-D), and primary MoDC experiments (Fig. 5B-E). Statistical comparisons in Fig. 4H-J were against no treatment.

For curli-producing versus curli-deficient EcN comparisons at each bacterial density (Fig. 3B), two-way ANOVA was used to assess the protein × concentration interaction, followed by planned two-sided Welch’s t-tests with Bonferroni correction at each concentration. For purified fiber dose-response data (Fig. 3D), two-way ANOVA was used to assess the protein × concentration interaction, followed by planned two-sided Welch’s t-tests with Bonferroni correction between engineered variants and wild-type curli at each concentration.

All experiments were performed with a minimum of n = 3 independent biological replicates unless otherwise specified in figure legends.

## DATA AVAILABILITY

All data supporting the findings of this study are contained within the manuscript and its supplementary materials. No large datasets, sequencing data, or computational models were generated or deposited in external repositories. Plasmid maps and relevant sequences are provided in the supplementary materials (Fig. S1, Table S1-2). Raw data for all figures are available from the corresponding author upon reasonable request.

## ACKNOWLEDGEMENTS

The authors thank William Fowle and Dr. Shirin Kaboli of the Boston Electron Microscopy Core of Northeastern University for technical assistance with TEM imaging. We appreciate the Institute for Chemical Imaging of Living System (CILS) of Northeastern University and Dr. Guo-Xin Rong’s assistance with Flow Cytometry. We thank Dr. Ke Zhang of the Department of Chemistry and Chemical Biology of Northeastern University for access to the Bio-Rad CFX96 Real-Time System for qPCR experiments. We thank Dr. Matthew Chapman and the members of the Chapman laboratory for their insights on curli purification. We are grateful to the Department of Chemistry at Tufts University for access to the J-1500 CD spectrophotometer. We thank STEMCELL Technologies for providing the human leukopak used in primary cell experiments. We thank Dr. Pedro Saavedra and Dr. Çağla Tükel for helpful discussions. We also thank members of the Joshi laboratory for critical feedback and discussion. Figures were created in BioRender. Joshi, N. (2026) https://BioRender.com/z1f6fkt.

Shanna Bonanno: Conceptualization, Data Curation, Formal Analysis, Investigation, Methodology, Project Administration, Resources, Supervision, Validation, Visualization, Writing – original draft, Writing – review & editing. Rutvi Sheta: Investigation, Resources, Validation. Teena Ramu: Investigation, Resources, Validation. Shriya Verenkar: Investigation, Resources, Validation. Darren Kim: Formal Analysis, Investigation. Eva Bessette: Formal Analysis, Investigation. Petit Pierre: Investigation, Writing – review & editing. Neel S. Joshi: Conceptualization, Funding Acquisition, Project Administration, Supervision, Writing – review & editing.

